# Single Cell Proteomics Reveals Novel Cell Phenotypes in Marfan Mouse Aneurysm

**DOI:** 10.1101/2025.02.15.638465

**Authors:** Louis Saddic, Giselle Kaneda, Amanda Momenzadeh, Lior Zilberberg, Yang Song, Mitra Mastali, Simion Kreimer, Alexandre Hutton, Ali Haghani, Jesse Meyer, Sarah Parker

## Abstract

**Background:** Single-cell omics technology is a powerful tool in biomedical research. However, single cell proteomics has lagged due to an inability to amplify peptides in a similar fashion to nucleotide strings. Single cell proteomics is important because proteins are the main functional unit in cells, and they often poorly correlate with mRNA quantities. In this paper we describe the first single cell proteomic analysis of complex tissue, comparing aneurysmal and normal mouse aorta from males and females. We also compare and integrate our single cell proteomic profiles with a matching single cell transcriptomics dataset.

**Methods:** We compared single cell proteomes between male and female, wild-type and *Fbn1^C1041G/+^* Marfan mice (N=3 per group). Individual cells from mouse aortic root single cell suspensions were deposited in 384 well plates and subjected to ultra-sensitive nanoflow liquid chromatography-ion mobility-time of flight-mass spectrometry. The data were analyzed with leiden clustering to identify cell types. Statistical analyses were performed to detect differential proteins within cell types and multi-omics analysis integrated single cell proteomics with published single cell RNA-seq.

**Results:** We identified all major aortic cell types including 7 distinct smooth muscle cell subtypes. The proportion of these cells varied based on sex and the *Fbn1^C1041G/+^* genotype. Differentially expressed proteins between male and female in addition to wild-type and Marfan samples uncovered enhanced endothelial to mesenchymal transition patterns in endothelial cells from male Marfan mice. Comparisons between single cell RNA and single cell proteomic profiles showed similarities in major subtypes but not smooth muscle cell subtypes. Multi-omics analysis of these two single cell platforms demonstrated a potential novel role for smooth muscle cell derived angiotensin signaling in the Marfan phenotype.

**Conclusions:** Single cell proteomics identified new subpopulations of vascular smooth muscles cells and novel cell type specific protein signatures related to sex differences and aneurysm formation.

**Abbreviations:** Next generation sequencing (NGS), Mass spectrometer (MS), Single cell proteomics by Mass Spectrometry (ScOPE-MS), Marfan’s syndrome (MFS), Fibrillin 1 (FBN1), Transforming growth factor β (TGFβ), Smooth muscle cell (SMC), Single cell proteomic (scProteomic), Differentially expressed proteins (DEPs), Wild-type (WT), Hanks’ balanced salt solution (HBSS), Fetal bovine serum (FBS), Dulbecco’s Modified Eagle Medium (DMEM), Data-independent acquisition parallel accumulation-serial fragmentation (DIA-PASEF), Magnetic assisted cell sorted (MACS), Single Cell Analysis in Python (Scanpy), Kyoto Encyclopedia of Genes and Genomes (KEGG), Principal component analysis (PCA), Uniform manifold projection (UMAP), Single cell transcriptomic (scTranscriptomic), Smoothelin (Smtn), Transgelin (Tagln), Myosin heavy chain 11 (Myh11), Platelet endothelial cell adhesion molecule 1 (Pecam1), Dipeptidase 1 (Dpep1), Uncoupling protein 1 (Ucp1), Low-density lipoprotein receptor-related protein (Lrp1), DNA ligase 3 (Lig3), Capsaicin channel transient receptor potential vanilloid 1 (Trpv1), Endothelial to mesenchymal transition (endMT), Intercellular adhesion molecule 1 (Icam1), Intercellular adhesion molecule 2 (Icam2), Endothelial cell-selective adhesion molecule (Esam), Calponin 1 (Cnn1), Vimentin (Vim), Zinc finger E-box-binding homeobox 1 (Zeb1), Snail family transcriptional repressor 1 (Snai1), Tropomyosin alpha-4 chain (Tpm4), Angiotensin converting enzyme (Ace)

## Introduction

The development of single cell RNA-seq was one of the most consequential advancements to the field of genomics and next generation sequencing (NGS). It allows un-biased characterization of individual cells using transcriptomic profiles in a way that has broadened our understanding of tissue diversity and cellular plasticity^1^. Transcripts are often taken as a proxy for protein abundance and expression, but due to numerous levels of biological regulation (e.g., RNA stability, protein translation rates, protein degradation rates), the correlation between RNA and protein expression can be variable from one gene to the next^2–5^. In addition, protein gene products can be extensively diversified through post-translational modifications which cannot be accurately inferred by RNA profiling^6^. Therefore, there is a strong interest in developing single cell proteomic technology to complement and extend cell phenotype analysis gleaned from transcriptomic profiling. Technical challenges around protein dynamic range, instrument sensitivity, speed and sample preparation, left single cell proteomic technology to lag somewhat behind RNA technology. Obstacles such as protein loss during sample processing and insufficient sensitivity in mass spectrometer (MS) instruments coupled with the inability to amplify peptides in a similar fashion to nucleic acid polymers makes it difficult to fully develop single cell proteomics to a comparable level relative to nucleic acid sequencing^7^.

Developments in single cell proteomics have been steadily advancing over the past several years. The very early applications reported single cell profiling of very large xenopus embryo cells, each with adequate protein content for detection on earlier generation mass spectrometers^8^. A leap in capability occurred with the development of the single cell proteomics by Mass Spectrometry (ScOPE-MS) approach, which uses tandem-mass-tag peptide labeling of individual cell peptides and multiplexes several cells into a single MS run, all alongside a ‘carrier’ channel of labeled peptides from a representative bulk lysate. While this enabled Bonafide single cell profiling, issues related to ion suppression and quantification have been challenging for the label-based approaches^9^. One consideration for highly heterogenous samples pertains to the likelihood of identifying a cell-specific protein from a rare cell type within a multiplex, when the selection of peptides for sequencing depends on amplification of numerous observations of a peptide across the cell types profiled and/or from the bulk carrier channel. This issue is overcome by direct label-free single cell proteomics analysis, which was recently made possible through a series of engineering advancements leading to the design of an ultra-sensitive mass spectrometer, the Bruker TIMSTOF SCP instrument^10^. Alongside advancements in the MS sensitivity, others have optimized nano-scale sample preparation set ups for minimal protein loss during tryptic digestion to further improve single cell profiling capabilities, and our group and others have advanced liquid chromatography set ups to enable fast, sensitive separation of peptides inline with MS to improve the throughput (e.g., number of cells profiled per 24 hours)^11^. All together, these advancements have launched LC MS based proteomics onto the map of options for single cell profiling, with a steady release of recent publications highlighting novel and complementary discoveries made by this technology^12^. To date, nearly all single cell proteomic studies published have focused on *in vitro* cell systems with moderate heterogeneity, raising the question of how the technology and data analysis approaches will perform on cellularly-complex organ systems.

Marfan’s syndrome (MFS) is an autosomal dominant genetic disorder resulting from mutations in the Fibrillin 1 (*FBN1*) gene. Patients develop phenotypic alterations in many organ systems but perhaps the most deleterious is aneurysms of the aortic root that carry a high risk of dissection and sudden death. In these patients, there is aberrant destruction of the medial layer of arteries leading to aneurysm growth of the aortic root and ascending aorta. Despite decades of knowing and researching the causal mutation driving MFS aneurysm, this has not translated to an impactful drug treatment to slow or stop aneurysm progression and as of now, the only treatment to Marfan aortic root aneurysms is high risk cardiac surgery^13, 14^. Mouse models of MFS, including the *Fbn1^C1041G/+^* mouse, recapitulate the aortic disease found in humans, and have been invaluable to uncovering the molecular causes that lead to this disease^15^. At first it was thought that excessive transforming growth factor β (TGFβ) signaling components in the media, due to failure of mutated Fbn1 to sequester the large latent TGFβ complex, causes vascular smooth muscle cell (SMC) mediated destruction of the extracellular matrix. Recent studies, however, using antibodies against TGFβ ligands or SMC specific knock-out of TGFβ signaling effectors, instead suggest that lack of TGFβ signaling to SMCs may also lead to aneurysm growth and dissection^16, 17^. Thus, there remains a considerable lack of clarity as to how MFS-causing Fbn1 mutations drive the molecular milieu causal of aneurysm progression and dissection, and it is likely that mapping cell-type specific responses to the Fbn1 mutation within the *in vivo* system are needed to compile a clear understanding of molecular pathogenesis. For instance, work in a related hereditary aortopathy, Loeys-Dietz syndrome, has shown that the embryological origins of the aortic SMCs result in differential TGFβ signaling sensitivities, subsequently driving SMC-subtype dependent responses to TGFβR mutations^18^. The critical role of SMCs in the pathogenesis of this disease is incontrovertible. In fact, there is significant evidence that the plasticity of SMCs allows them to shift their phenotype into a diversity of cell types that ultimately unravels the architecture of the aortic wall. It is also clear that other vascular cells, including endothelial and immune types, are altered in MFS aneurysm and likely play a role in the initiation and/or progression of aneurysms, or the final failure of wall integrity leading to dissection or rupture^19, 20^. Identifying the integrated response of cells to the Fbn1 mutation, and the causes and identity of cellular phenotype shifts could be the key to reversing disease progression and perhaps open the door to novel therapeutics.

Our group recently developed a workflow for high-throughput direct single cell profiling of up to 96 cells per day, which enabled proof of concept single cell proteomic (scProteomic) profiling on one mouse aorta, representing the first complex tissue to be analyzed by scProteomics^11^. Vascular tissue is an ideal system to test this new technology as it is not only comprised of a diversity of different major cell types (e.g., endothelial, SMC, fibroblast, immune) but also because the most abundant cell type, the SMC, is well known for its high degree of plasticity and ability to adopt a diversity of fates and sub-phenotypes. Here we show that single cell proteomics can accurately profile individual cell types in mouse aortic root tissue and further subdivide SMCs into distinct subgroup populations. In addition, we show that this new technology can uncover differences in the abundance of these cells and cell-specific differentially expressed proteins (DEPs) based on sex differences and the Marfan phenotype. Even more, we show that combining single cell RNA with single cell proteomic profiles of Marfan mouse aortic roots can identify novel pathways that may play a role in aneurysm biology.

## Methods

### Isolation of Mouse Aortic Root Cells and Processing for Proteomic Analysis

All procedures involving mice were approved by the Cedars-Sinai IACUC review panel. A total of N=24 male and female, Wild-type (WT) and Marfan *Fbn1^C1041G^* mice were sacrificed between 12-13 weeks of age using high concentration isoflurane inhalation. The inferior vena cava of the mouse was cut to let blood flow out and 10 ml of ice-cold PBS was injected into the left ventricle. Aortic roots from the aortic valve to 1-2mm above the sinotubular junction were dissected and cleaned of most valvular, myocardial, other connective tissue and excess/loosely adhering pericardial fat tissue. Aortic roots from N=2 mice of matching genotype and sex were rinsed of blood with ice-cold PBS and then transferred in prewarmed enzymatic digestion cocktail prepared with 300U/ml of collagenase Type II (Worthington CLS-2), and 3 U/ml of elastase (Worthington ES) in Hanks’ balanced salt solution (HBSS). Aortas were diced into small pieces and incubated at 37C for 40 min to allow dissociation of cells from the connective extracellular matrix. The digestion solution was gently passed 10x through a 21-gauge needle to further liberate remaining cells and enzymatic reaction was stopped with 20% fetal bovine serum (FBS) in Dulbecco’s Modified Eagle Medium (DMEM). The mixture was then passed over a 70um cellular strainer to collect dissociated cells and remove matrix tissue and cell clumps. The strainer/filter was washed with ice-cold PBS and cells suspension was centrifuge at 500g at 4C for 5min. Cell pellets containing approximately 3 – 5 x 10^4^ cells were resuspended in 200ul ice-cold PBS. Cell suspensions were then labeled with Sytox Green nuclei acid stain. Specifically, 1ul of 5mM Sytox green solution was added to the 500uL cell suspension (0.5×10^6^cells/mL) and incubated at room temperature for 15 minutes. The cell suspensions were then transferred into the loading vial of a Scienone CellenONE cell sorting and robotic liquid dispensing instrument. Cells ranging in diameter from 10 – 40uM with minimal green fluorescence (e.g., live cells) were dispensed one per well of a 384 well plate that was preloaded with 200nL of lysis buffer (100mM TEAB, 0.2%DDM and 100ng/uL trypsin in 50mM acetic acid). Plates of sorted, lysed cells were stored no more than 60 days, covered at –80C until further processing. Lysates were digested by dispensing 200nL of 40ug/mL trypsin into each plate well, and incubated for 4-hours at 37C before quenching digestion with 200nL of 0.1% Formic Acid solution. Quenched digested peptide was dried completely, and plates were placed on an autosampler for liquid chromatography mass spectrometry analysis.

### Direct Single Cell Proteomic Data Acquisition

LC-MS was based on the nano dual trap single column (nanoDTSC) method described in Kreimer et al^11^. Briefly, the contents of each well on the 384-well plate were resuspended in 20uL of 0.1% formic acid 2% acetonitrile using the autosampler and injected onto one trapping column (Exp2 170nL packed with 10um diameter PLRP beads, Optimize Technologies). Concurrently the previous sample trapped on an identical trapping column was eluted onto the analytical column (15cm x 75um filled with 1.9um particles PepSep Bruker) and separated using a binary reversed phase gradient (0.1% formic acid in water to 0.1% formic acid in 80% acetonitrile 20% methanol) at 500nL/min flowrate. The gradient was delivered as follows: 9%B to 25%B over 8 minutes 25%B to 38%B over 4.6 min, followed by a 98%B wash at 98%B at 1000nL/min for 1.2 min and equilibration to 9%B at 100nL/min for 1 min (15 min total run time or 96 samples/day). The separated peptides were sprayed through a 20um ZDV emitter installed in the Bruker captive source at 1700V with the dry gas set at 3.0L/min and 200C glass capillary temperature. Data were acquired by data-independent acquisition parallel accumulation-serial fragmentation (DIA-PASEF) with the ion accumulation and trapped ion mobility ramp set to 166 ms. Each MS1 scan was followed by 90m/z wide DIA scans spanning 300– 1200m/z and 0.6–1.43 1/K0 with 4 trapped ion mobility ramps (0.86s total cycle time).

### Generation of the Sample-Specific Peptide Library

The final peptide assay library used to search single cell files was produced by merging three separate libraries: Bulk DDA, LibraryFree DIANN selected single cell, and a preliminary library produced from our prior publication^11^. Methods for each are as follows and summarized in Supplementary Figure 1 A-E.

*Bulk DDA library.* Magnetic assisted cell sorted (MACS) CD31+ and CD31-fractions of dissociated mouse aortic root cells as well as MACS sorted CD45+ peripheral blood mononuclear cells, all collected from a WT and MFS mouse, were run separately by cell source / genotype (e.g., N=6 final DDA samples run). Isolated cell pellets were lysed using 8M Urea/5% SDS and processed for tryptic digestion using standard procedures except that no reduction of cysteine residues (e.g., no DTT or IAA alkylation step) was performed in order to produce peptides consistent with the single cell sample processing protocol. Protein lysates were processed using the STRAP manufacturer recommended protocol (Protifi, Fairport, NY). Peptides were analyzed on the Bruker TIMSTOF SCP instrument using the same LC configuration but with data dependent acquisition (PASEF-DDA). Both accumulation and ramp times were set to 166 ms with DDA scans acquired over 5 ramps within a polygon spanning 240-1700m/z and 0.6-1.5 1/K0 for a cycle time of 1.03 s. The resulting raw files were searched using the FragPipe MS Fragger workflow^21^ and integrated into a ezPQP library.

*LibraryFree DIANN selected single cell.* The full dataset of single cell files were initially searched against our previously published preliminary library. The resulting protein matrix was analyzed using the Single Cell Analysis in Python (Scanpy)^22^ workflow, as described below, and preliminary cell type assignments were made according to leiden clustering and canonical cell type markers. From these preliminary cell type assignments, the single cell data files with the top 10 highest protein identification counts from each cell category were subsequently run in DIANN using a library free search. The product of the library free search includes a spectral library. These sample-specific spectral libraries from each searched cell type category (SMCs, endothelial cells, adipocytes, fibroblasts, macrophages/immune cells, mesothelial/epithelial cells) were sequentially merged, by appending only unique peptides not seen in any of the other libraries.

*Final Merged Library.* The final library was produced using a sequential merge in which first only peptides unique to the library free search were appended to the Bulk DDA library. Subsequently, only unique peptides from the preliminary library were added to the previously merged library of Bulk DDA and Library Free search results. The final library contained 45,287 peptide precursors and 5,345 protein groups and was used for all single cell analyses.

### Peptide Sequence Identification and Protein Quantification Estimation

Raw files were searched using DIANN v1.8.1^23^ against the custom assembled sample specific library (see above), using matches between runs and reannotation of gene groups to a supplied mouse FASTA protein sequence database, but with no heuristic protein inference and no normalization applied. Protein quantification estimates were produced using the DIANN maxLFQ algorithm, and only proteins quantified by proteotypic peptides were included in the final dataset for further analysis.

### Bioinformatic Analysis of Single Cell Proteomic Data

Data QC and analysis were performed using the Scanpy python package^22^ The final protein-by-cell data matrix was uploaded and converted into an annotated data matrix object in the Scanpy single cell data analysis workflow^22^. Cells with fewer than 200 proteins identified and proteins observed in fewer than N=3 cells were dropped before normalizing individual intensities to the total signal from each cell, scaling by log10 and performing a regression adjustment on total protein counts observed for each cell. Batch effect correction was performed using ComBat^24^, with each plate considered a separate batch. Batch corrected protein matrices were then processed using PCA, leiden based clustering (using default parameters except resolution was set to 1.2), and UMAP dimensionality reduction and visualization. General cell type annotation was deciphered using canonical marker proteins, with unbiased marker analysis performed for corroboration as well as identification of cell subtype markers. Cell proportions were compared between groups at the level of all observed cells per group, and all cells observed per biological replicate. Differential protein analysis within cell type groups was performed using linear mixed effect modeling, mandating that a protein was seen in at least N=3 cells of at least one comparison group within each cell type. Figures were generated using Scanpy or after exporting the Scanpy annotated data matrix object and conversion into Seurat for figure generation and analysis in R^25^.

### Comparison of Single Cell Proteomic Data to a Similar Published Single Cell RNA Sequencing dataset

Raw counts were downloaded from the Gene Expression Omnibus database using the identifier GSE186845 belonging to the datasets described in Pedroza et. Al. from 16-week-old wild-type and MFS *Fbn1^C1041G^* male and female mice (N=1 per group)^19^. Single cell transcriptomic (scTranscriptomic) data were re-analyzed using the same Scanpy workflow used to analyze the protein data, with the exception that batch effect correction was not performed. Marker genes either matching the canonical scProteomic cell type marker set or derived from the original source publication were used to ascertain cell-type cluster identifications, with unbiased marker analysis also being performed. The scTranscriptomic and scProteomic data objects were then exported and converted to separate Suerat objects in R using the Schard package (https://github.com/cellgeni/schard). An initial comparison of the two datasets was performed using the Seurat package ‘FindTranferAnchors’ and ‘TransferData’ functionality^26^, in which shared marker proteins from one data object are used to assign cell types to the other dataset for comparison of how reannotated cell identities cluster together relative to their original annotations. We then used the LIGER^27^ multi-omic integration package in CRAN^28^ to merge the count matrices from the proteomic and RNA datasets, with k=10 and lambda=5 used for the ‘runIntegration’ merging step. The merged LIGER object was quantile normalized after which leiden clustering (nNeighbors =30, resolution=0.5) and UMAP dimensionality reduction (nNeighbors=30, minDist=0.3) were performed. Canonical marker genes/proteins were used to estimate cell type annotations onto new, integrated LIGER leiden clusters. A Sankey plot was constructed comparing the groupings of any given cell’s annotation between their parental dataset (e.g., RNA or protein only) and the integrated, LIGER dataset. The proportion of cells from each genotype, whether from RNA or protein datasets, were calculated across the new LIGER clusters. To assess whether the proportions of sex (male vs. female) and genotype (MFS vs. WT) differed significantly across clusters, we performed two-sided z-tests for proportions. First, the dataset was reformatted into a long format, where each observation contained the LIGER cluster assignment, sex, genotype, and count. We computed the overall proportions of females and MFS samples across all clusters as reference values. For each cluster, we compared its sex-specific and genotype-specific proportions to the overall dataset proportions using a z-test for two proportions, which tests whether the proportion in a specific cluster significantly deviates from the expected proportion under the null hypothesis of no difference. Finally, we analyzed marker features (shared protein and transcript observations) across the LIGER leiden clusters using its built-in Wilcoxon test of expression in the test cluster relative to all other clusters. Kyoto Encyclopedia of Genes and Genomes (KEGG) pathway enrichment^29^ of the top 100 shared marker features with Log_2_FC > 1 for a given cluster expression was analyzed in the ENRICHR platform^30^.

## Data availability

Raw data, processed counts matrix, and the Scanpy and Seurat objects used to generate these results are deposited on the MassIVE server for public access.

## Results

### Single Cell Proteomics identifies major aortic cell subtypes

Wild-type and *Fbn1^C1041G/+^* Marfan mice at 12-13 weeks of age were selected for single cell proteomic analysis in this study. Marfan mice had significant dilation of the aortic root visible upon gross dissection compared to wild-type littermates. Two aortic root cell suspensions per biological replicate were pooled to yield sufficient cells for downstream sorting. Within each genotype, three biological replicates for each sex were selected bringing the total sample size to 24. Mouse aortic tissue was taken from above the aortic valve to 1-2mm above the sinotubular junction and subjected to enzymatic digestion to generate single cell suspensions. Individual cells were then sorted and deposited onto a 384-well plate using the CellenONE system followed by lysis and tryptic protein digestion. Single cell proteomes were then acquired on the Bruker TIMSTOF SCP mass spectrometer using the nonDTSC workflow (Figure 1A). A total of 2,746 protein groups were detected from 157,743 peptides across 4,986 cells. We eliminated cells with fewer than 200 proteins as well as proteins not identified by unique, unambiguous peptide sequences or detected in less than three cells during quality control filtering. This resulted in a total of 3,475 cells and 2,421 proteotypic proteins available for downstream analysis, with an average of 487 identified proteins per cell.

**Figure 1.**
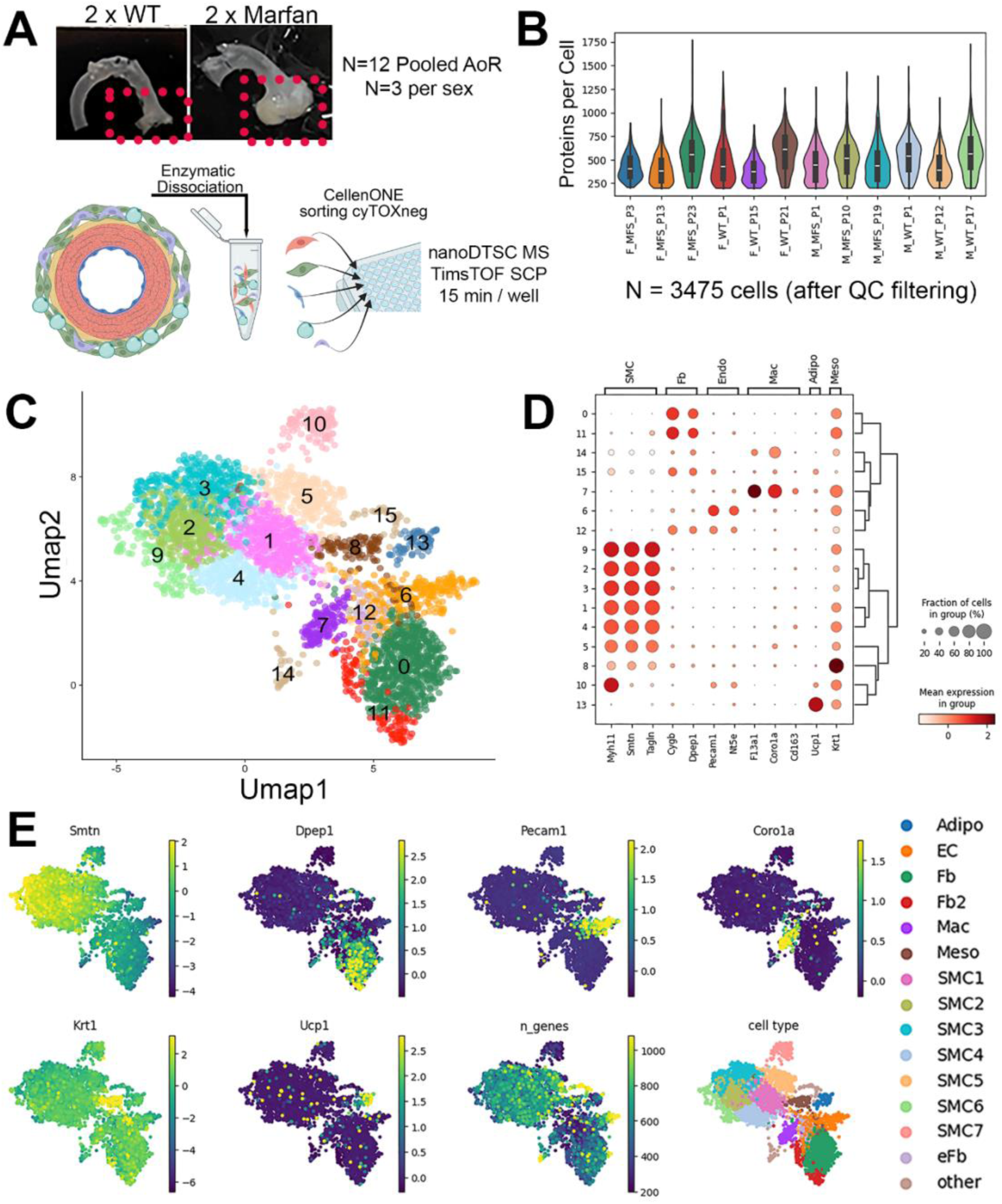
Experimental workflow and results overview. (A) Overview of the experimental workflow. Aorta (top panel) from wild type (WT) and *Fbn1^C1041G^* Marfan’s syndrome (MFS) mice were enzymatically dissociated to release individual cells, which were subsequently sorted on a CellenONE system into 384-well plates for lysis and tryptic digestion ahead of acquisition of individual cell proteomes on the Bruker TIMSTOF SCP mass spectrometer using the nanoDTSC workflow. (B) Distribution of proteins identified per cell, separated by processing batch. (C) Projection of all mouse aortic cells onto UMAP dimensionality reduced space, with 16 sets of proteomically-similar cells (e.g., cell types) mapped as separate colors using leiden algorithm. (D) Dot plot to display the expression percentage and average intensity of canonical cell type markers for known aortic cells across the 16 identified leiden clusters. (E) Cell phenotype assignments based on canonical expression patterns.

Of note, these counts and protein identification ranges were distributed evenly across biological replicates and plates (Figure 1B and Supplementary Table 1). After initial dimensionality reduction and clustering, we observed uneven mixing of batches and biological replicates across clusters (Supplementary Figure 2A), prompting batch correction using ComBat^24^ correcting on cell processing plate ID. This ensured that all biological replicates were well balanced across most cell clusters (Supplementary Figure 2B).

We first jointly analyzed all 3,475 cells from all samples (male/female and wild-type/Marfan). Using Scanpy, leiden clustering based on neighbors computed from principal components discovered 16 groups. These clusters were then visualized in reduced space determined by UMAP (Figure 1C). Analysis of specific marker genes for aortic cell types including smoothelin (Smtn), transgelin (Tagln), and myosin heavy chain 11 (Myh11) for SMCs, platelet endothelial cell adhesion molecule 1 (Pecam1) for endothelial cells, dipeptidase 1 (Dpep1) for fibroblasts, and uncoupling protein 1 (Ucp1) for adipocytes (Figure 1D and E), indicated that these cell groups included 7 subtypes of smooth muscle cells, 2 subtypes of fibroblasts, endothelial cells, adipocytes, macrophages, and mesothelial cells. There were an additional 2 clusters that could not be clearly identified and are annotated as ‘other’. The top 15 marker proteins identified for each cluster are plotted in Supplementary Figure 3 and marker DEP tables from Scanpy are provided in Supplementary Table 2.

### Unique proteome signatures define smooth muscle cell subtypes

Not surprisingly, SMCs were the predominant cell type from aortic samples. As mentioned previously, leiden clustering of cells from all samples identified 7 distinct subtypes of SMCs (Figure 2A). When comparing the global proteome of all cell types in the aorta, all SMC subtypes were most similar to other SMC subtypes with the exception of SMC subtype 7 which shared more similarity with adipocytes (Figure 2B). SMC7 was designated an SMC due to its high abundance of Myh11 despite low expression of other SMC markers such as Smtn and Tagln. In fact, SMC7 had the second highest intensity of Myh11 compared to all other SMC subtypes (Figure 2C and 2D). When we examined some of the protein abundance patterns among the different subtypes of SMCs a few relationships emerged. SMC2, SMC3, and SM6 were in general classified as having high abundance of the known canonical SMC markers namely Myh11, Smtn, Tagln, and vinculin (Vcl). On the other hand, SMC1, SMC4, and SMC5 had slightly lower abundance of these canonical markers of SMCs and higher abundance of the proteins low-density lipoprotein receptor-related protein (Lrp1) and DNA ligase 3 (Lig3). SMC1 and SMC5 also had high abundance of protease serine 2 (Prss2), and SMC5 had uniquely high abundance of the capsaicin channel transient receptor potential vanilloid 1 (Trpv1) (Figure 2C and 2D). Given the relatively lower abundance of canonical contractile markers in the SMC1, SMC4, and SMC5 subtypes, they may represent phenotypically modified smooth muscle cells with more synthetic like functions compared to the classical contractile smooth muscle cells represented by clusters SMC2, SMC3, and SMC6.

**Figure 2.**
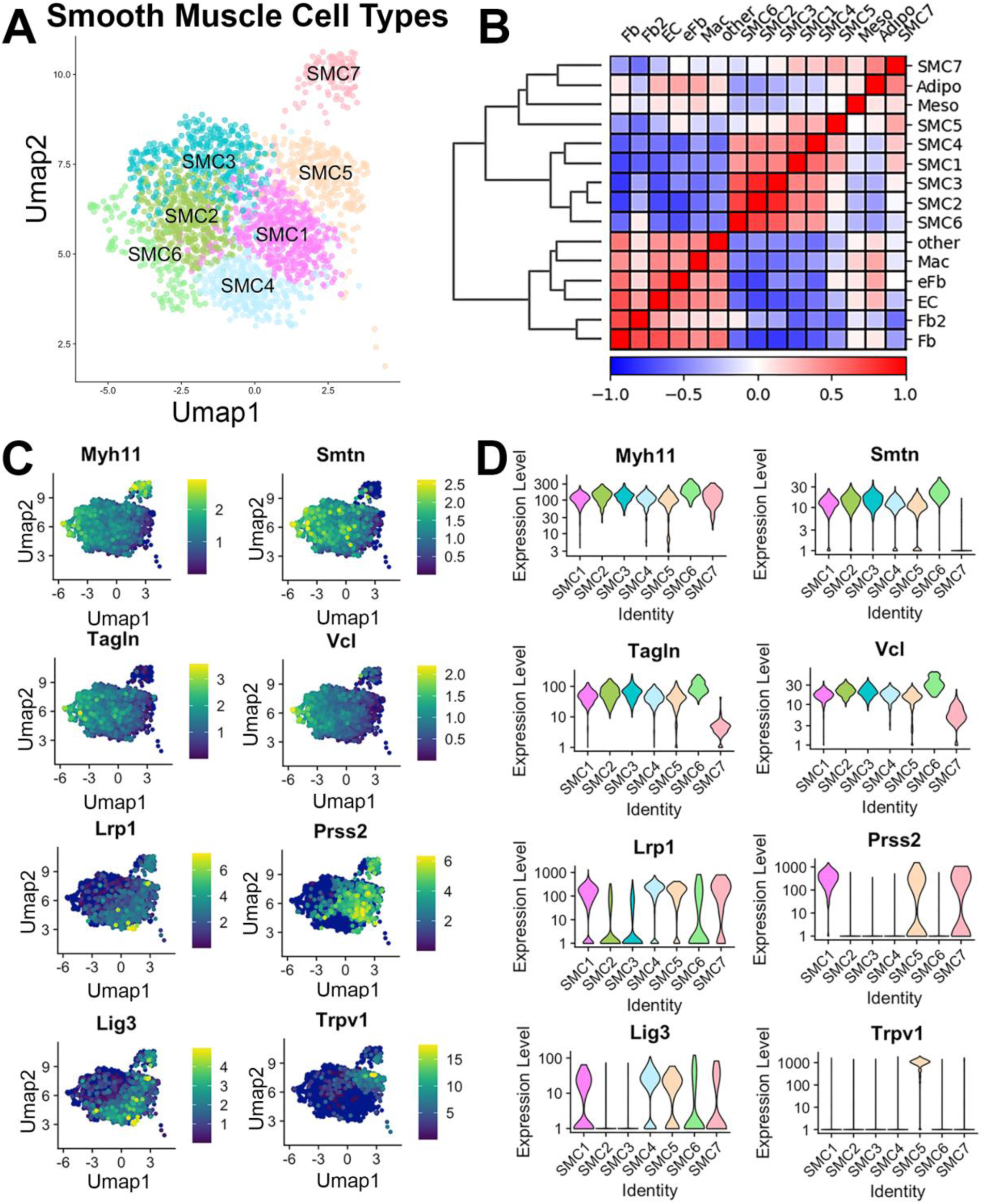
Deeper analysis of Smooth Muscle Cell (SMC) Sub-phenotypes. (A) UMAP projection of only SMC – cell sub phenotypes (SMC1-SMC7). (B) Correlogram of proteomic similarity between and across all cell types identified. (C) Projection of identified SMC – subtype marker proteins. (D) Violin plots of quantified marker expression levels across SMC sub-phenotypes.

### Sex and Genotype alter abundance of cell types

Cells for biological replicates (n=3) were pooled for each sex within each genotype and the corresponding UMAPs and relative abundance of each cell type within each sex and genotype category were plotted (Figures 3A-C and Supplementary Table 3). Male mice exhibited a higher relative proportion of endothelial cells (10.0% vs 4.9% of all cells, p-value 0.02), macrophages (5.3% vs 2.4%, p-value 0.023) and fibroblasts (21.7% vs 14.8%, p-value 0.0049) compared to female mice regardless of genotype. Marfan mice, on the other hand, showed most notable cell proportion differences within the Male mice, which had significantly fewer of the quiescent SMC3 cell type (12.5% of all cells in WT vs 4.2% in MFS, p-value 0.04) and significantly more of the mesothelial cell type (4.2% of all cells in MFS vs 2.1% in WT, p-value 0.02), and trended toward relatively higher proportions of macrophages (7% in MFS vs 3.6% in WT, p-value 0.1) and modified SMC cluster SMC7 (6.8% in MFS vs 3.1% in WT, p-value 0.17).

**Figure 3.**
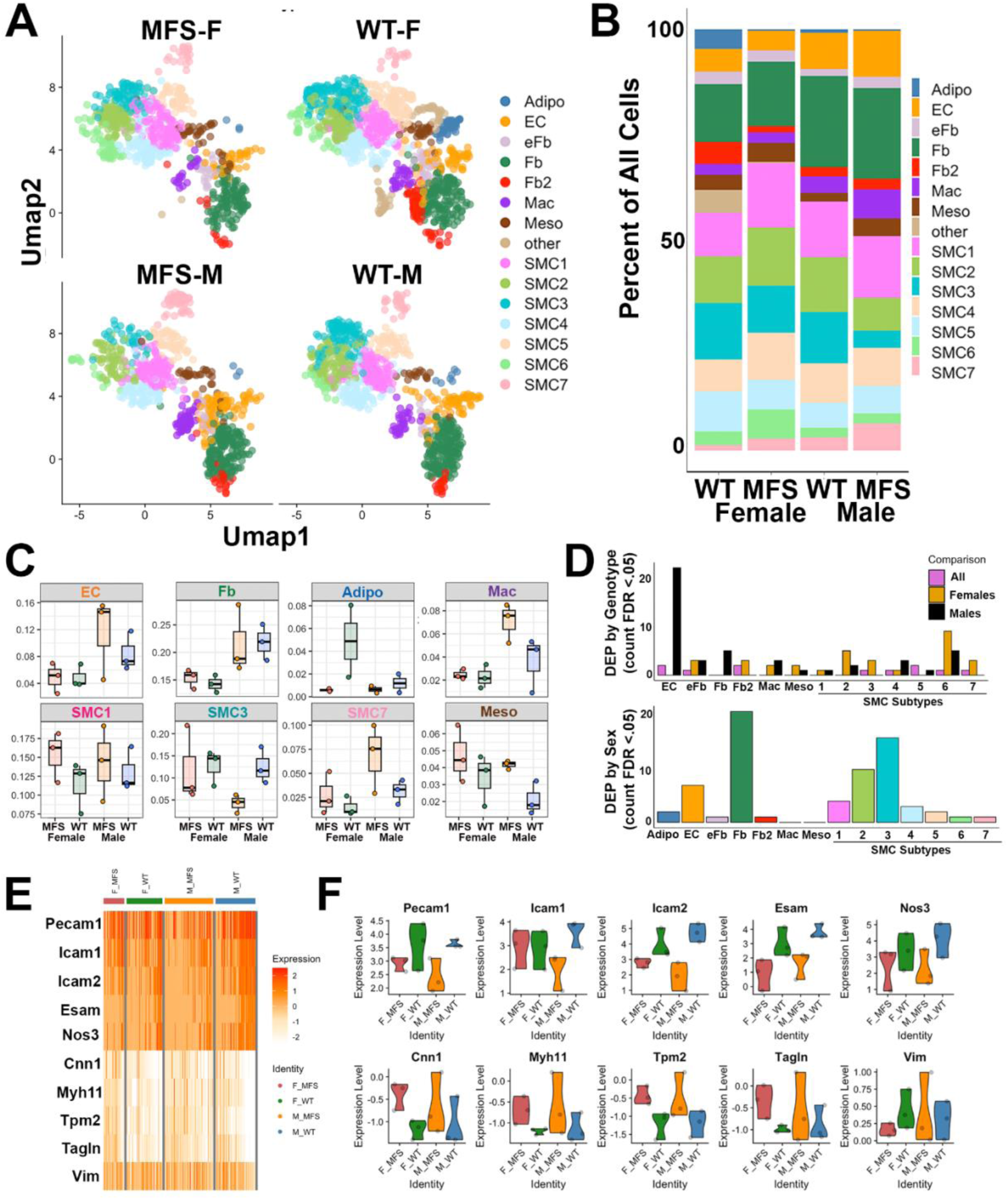
Comparison of Sex and Genotype patterns across single cell proteomic data. (A) UMAP projection of aortic cell types separated by donor mouse sex and genotype. (B) Relative proportion of each assigned cell type across all cells identified per mouse sex and genotype category. (C) Selected proportions of a selected cell type based on percentage assigned within an individual mouse biological replicate (N=3 biological replicates per sex and genotype combination). (D) Count of significant differentially expressed proteins (FDR < 0.05) from linear mixed effects modeling comparing genotype (upper panel) or sex (lower panel) within each assigned cell type. For genotype, effect of both sexes combined was compared to differentially expressed proteins (DEPs) within males (M) and females (F) (by genotype) separately. (E) Subset of key proteins potentially indicative of endothelial to mesenchymal (endMT) transition with expression intensity in Marfan’s syndrome (MFS) vs wild-type (WT) endothelial cells. (F) Violin plots of endMT indicators across male and female MFS and WT endothelial cells.

We also compared the number of DEPs between wild-type and Marfan mice and between male and female mice within each cell type using linear mixed effects modeling and an FDR cutoff of 0.05 (Supplementary Tables 4, 5, and 6). Overall, there were relatively few proteins in each cluster that met false discovery rate cutoff (<0.05), with larger numbers of possible DEPs apparent by nominal p-values (Supplementary Figure 4A). There were higher numbers of sex specific DEPs in fibroblasts, endothelial cells, and certain SMC subtypes. This could partially be explained by higher cell abundances in these cell types but also may represent sex-specific gene regulation differences.

Within genotype, we combined males and females and performed the analysis between wild-type and Marfan, and then we repeated the comparison of genotype within cells of each sex separately. As with the sex DEPs, a relatively small number of proteins met FDR cutoffs to be confidently considered DEPs, with larger numbers of putative DEPs possible according to nominal p-values (Supplementary Figure 4B). Interestingly, there were noticeably more differentially expressed proteins by genotype in endothelial cells of male mice, but few of these proteins were also significantly differentially expressed in female mice or when males and females were combined for genotype comparison (Figure 3D). When we further analyzed the identity of the DEPs in male MFS vs WT endothelial cells, we noticed that they represented patterns that could be associated with the process of endothelial to mesenchymal transition (endMT). Compared to female and wild-type mice, Male Marfan mice had lower expression of endothelial adhesion proteins such as Pecam1, intercellular adhesion molecule 1 (Icam1), intercellular adhesion molecule 2 (Icam2) and endothelial cell-selective adhesion molecule (Esam) along with higher expression of smooth muscle proteins such as calponin 1 (Cnn1), Myh11, Tagln, and vimentin (Vim) indicative of a mesenchymal like state (Figure 3E and 3F). Interestingly, analysis of endothelial cell types profiled in previously published single cell transcriptomic data from *Fbn1^C1041G/+^* mice (further analysis details below) showed that mRNA for two transcriptional regulators of endMT, zinc finger E-box-binding homeobox 1 (Zeb1) and snail family transcriptional repressor 1 (Snai1), were both elevated in MFS endothelial cells relative to WT (Supplementary Figure 5).

### Comparison of singe cell RNA and Protein in Marfan mice uncovers key differences in cell type assignment

Previous studies have reported the single cell transcriptome of the *Fbn1^C1041G/+^* mouse model used in this study, enabling us to explore the degree of similarity between the two single cell modalities. Using the same Scanpy pipeline applied to our scProteomic results, we reanalyzed publicly available data from the Gene Expression Omnibus produced from single cell transcriptomic analysis of similarly aged *Fbn1^C1041G/+^* and wild-type mice, generating a UMAP sorted plot of single cell RNA-seq derived cell clusters and marker genes that were comparable to those produced by the original authors Pedroza et. al. (Supplementary Figure 6)^19^. Using Seurat anchor marker analysis, we compared cell marker annotation between our proteomics and the previously published Pedroza et. al. transcriptomic datasets^31^. In this analysis, trans-analyte plots in UMAP space were generated from cell type markers from RNA-seq to define cell types in the proteomic data and vice versa (Figure 4A, originally defined subtypes on left panels, projected trans-analyte defined cell types on right). While major subtypes were consistent in the trans-analyte analysis (e.g., SMCs, fibroblasts, macrophages, endothelial cells), cell subtypes, such as different categories of SMCs, were less preserved. We then performed an integrated multi-omic analysis using LIGER to combine the RNA and protein datasets into a single data matrix. Re-clustering using leiden (Figure 4B) with assigned cell types based on composite marker expression generated a new plot in UMAP space (Figure 4C). The expression level and location of several of these composite cell type markers demonstrate the specificity of these genes for defining the multi-omic clusters (Figure 4D). A Sankey plot was used to illustrate the contribution of each cluster from the RNA dataset and the protein dataset to the merged leiden clusters from the multi-omic LIGER algorithm (Figure 4E). As with the Seurat marker projection comparison, even with multi-omic integration the two datasets are only broadly matched by general cell types, wherein more subtle cell sub-phenotypes did not contribute in any clear patterns to the integrated clusters within the LIGER dataset.

**Figure 4.**
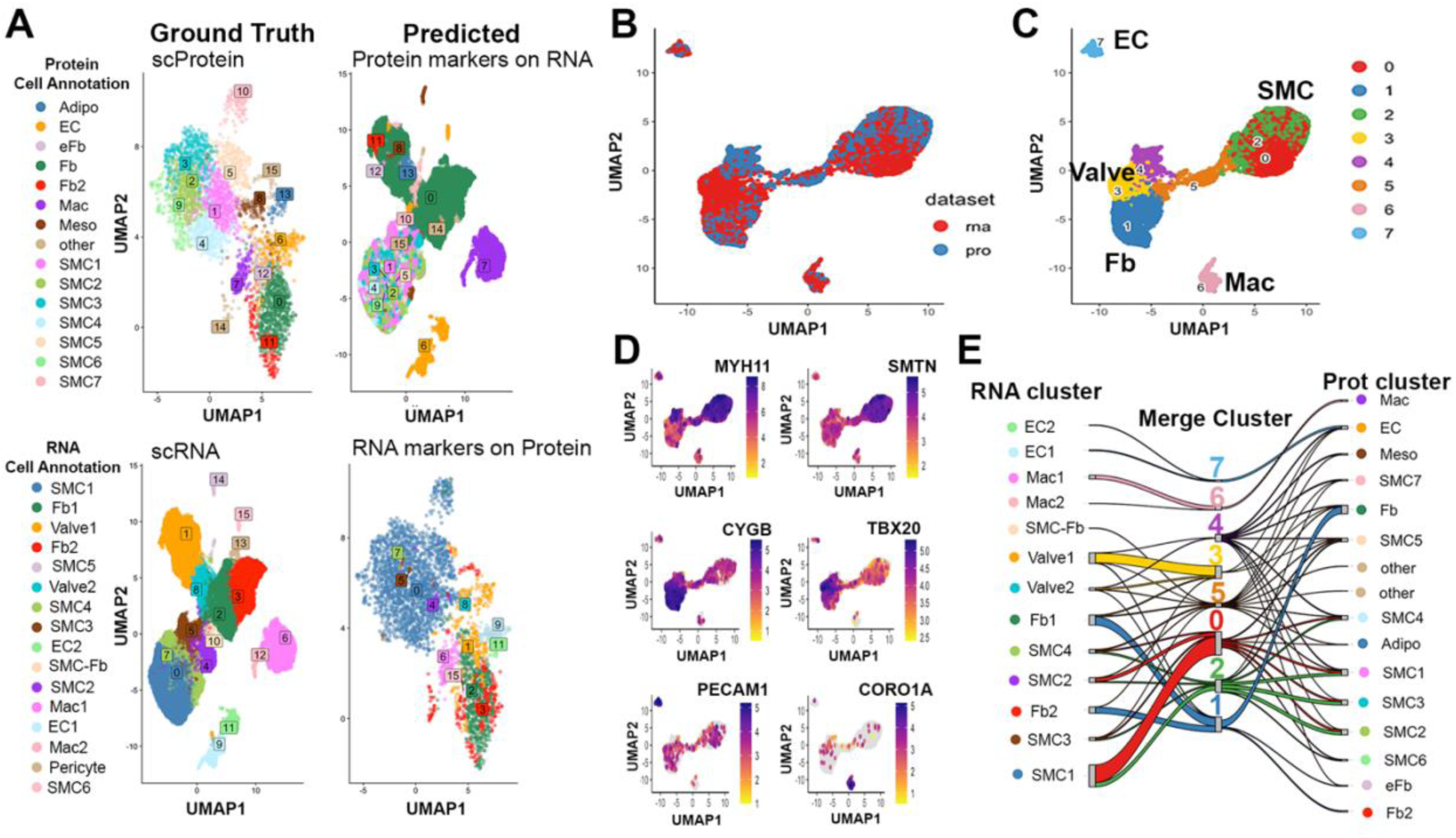
Comparison of scProteomic and scTranscriptomic sequencing data from *Fbn1^C1041G^* Marfan’s Syndrom (MFS) mice. (A) Seurat analysis of marker projections between transcriptomically defined and proteomically defined mouse aorta cell types. Left upper and lower panels represent the original cell types as defined by cis-analyte types (e.g., proteomic data defined by proteomic markers). Right upper and lower panels show cell types as defined by trans-analyte types (e.g., proteomic data defined by RNA markers). (B) Integrated multi-omic analysis using LIGER showing cells projected onto UMAP dimensionality reduced space, colored by dataset of origin (RNA vs proteomic). (C) Leiden clusters and corresponding assigned cell types on multi-omic datasets based on overall marker expression profiles. (D) Expression patterns of selected marker genes used to assign general cell types to multi-omic leiden clusters. (E) Sankey plot demonstrating differences in cell cluster assignment of a given cell from the transcriptomic (RNA clusters, left) or proteomic (protein clusters, right) datasets into their corresponding multi-omic defined leiden cluster from the LIGER integrated dataset (middle).

### Multi-omic analysis of SMCs uncovers unique gene signatures enriched in Marfan mice

To further examine whether similarities in cellular sub-phenotypes could be identified from the multi-omic analysis, we focused specifically on SMCs. To do this, we subset each dataset to only the cells annotated as SMC, resulting in 2200 total SMCs from the RNA dataset and 2031 total SMCs from the protein dataset. As with the full dataset, the two SMC datasets were integrated using LIGER, and a SMC-specific plot of cells in dimension reduced space derived from UMAP of integrated multi-omic data was generated (Figure 5A left), with 7 unique SMC subtype leiden clusters estimated (Figure 5A right). The proportional contribution of cells based on sex and genotype to each leiden cluster demonstrates significant enrichment in cluster 6 among mice known to typically have smaller aortic dimensions (7.5% in WT female, 4.1% in MFS female, 1.5% in MFS male and 2.7% in WT Male) and enrichment in clusters 1 and 4 in the MFS mice ( 22.8% vs 12.5% in MFS vs WT female and 26.3% vs 12% in MFS vs WT male for Cluster 1; 12.6% vs 8.7% in MFS vs WT female and 14.4% vs 8.0% in MFS vs WT male for Cluster 4) (Figure 5B, Supplementary Figure 7,and Supplementary Table 7). As with the full dataset, even with this focused SMC analysis, Sankey plot demonstrates relatively poor overlap between RNA defined SMC subtypes and protein defined SMC subtypes relative to how they contribute to the integrated multi-omic LIGER SMC clusters (Figure 5C). To understand more the biological similarities and differences between these multi-omic-defined clusters, we determined the top marker features (e.g., shared RNA transcript and protein between datasets) for each cluster (relative to all other clusters) (Figure 5D and Supplementary Table 8), and performed a KEGG pathway enrichment analysis (Figure 5E and Supplementary Tables9A-G). While canonical SMC markers did not meet the cutoff of ‘top marker’ for any of the clusters, there were noticeable differences between clusters, with the MFS enriched clusters 4 and 7 demonstrating lowest combined levels of most contractile markers (Supplementary Figure 8). In terms of top markers, two particularly interesting markers of MFS enriched cluster 4 were tropomyosin alpha-4 chain (Tpm4) and Ace, markers of MFS enriched cluster 1 included S100A4 and Annexin A1 and A3, and cluster 7, which trended toward upregulated in MFS, showed uniquely high Lrp1 among its top 5 differential features (relative to all other clusters). KEGG pathway analysis of expanded marker sets (up to 100 top significant markers with Log_2_FC >1) revealed that clusters 0, 5, and the WT enriched cluster 3 were particularly enriched in smooth muscle contraction features, while cluster 1 and 4 showed enrichment for actin cytoskeletal organization and ECM –receptor interaction, and cluster 4 was uniquely enriched in features related to the phagosome. In summary, use of the combined transcriptome and proteome signals differently defined subtypes of SMCs, and these multiomically defined subtypes revealed Marfan specific shifts in cell proportions, with alterations in contractile apparatus, cytoskeletal / ECM, phagocytosis and other biological pathways highlighting these MFS-specific changes in cell proportion.

**Figure 5.**
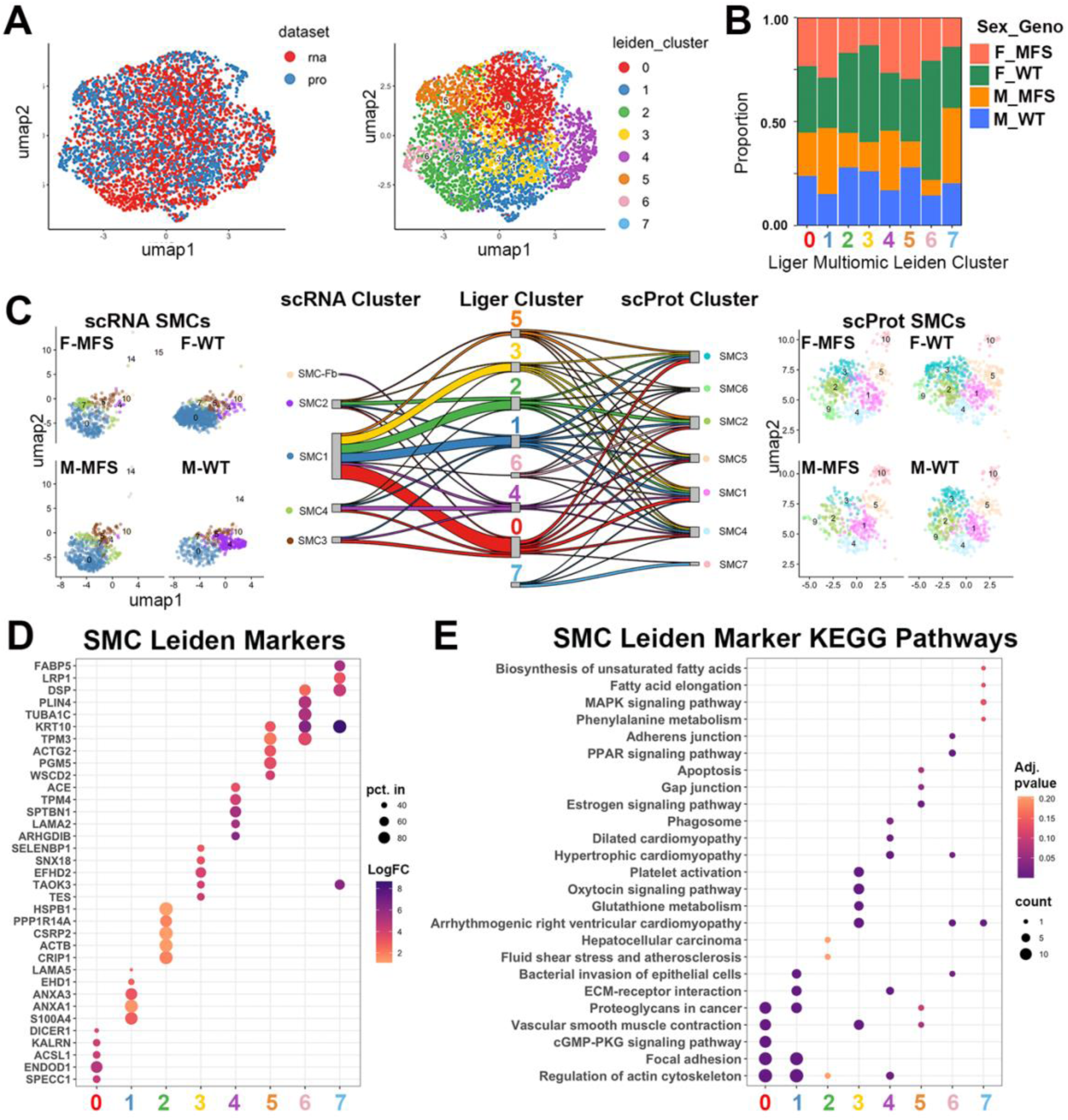
Focused Multi-omic analysis of smooth muscle cell (SMC) subtypes. (A) UMAP projections of SMCs as defined within respective RNA and proteomic datasets, demonstrating integrated mixing (left panel) and leiden SMC subtype clustering (right panel). (B)Proportion of multi-omic SMCs by mouse sex and gender separated across leiden defined SMC subtypes. (C)Sankey plot demonstrating mixing of cells from originating RNA-defined (left) and proteomic defined (right) clusters into multi-omic defined (middle) clusters. (D) Dot plot depicting the top shared marker features from the RNA and protein datasets for each LIGER derived leiden cluster. (E). Dot plot describing significantly enriched KEGG pathways from the shared marker features for each LIGER derived leiden cluster. M indicates male; F, female; WT, wild-type; and MFS, Marfan’s syndrome.

## Discussion

As the technology of next generation sequencing continues to expand exponentially, we grow increasingly close to mapping every macromolecule in every cell of a living organism. Multi-omics wouldn’t be complete without protein information, and in this manuscript, we describe one of the first examples of single cell proteomics by mass spectrometry applied to animal tissues. Mouse vasculature was chosen as a model tissue due to its assortment of well-defined cell types, and Marfan aneurysm presents an excellent test case as a hereditary condition driven by a monogenetic driver affecting an extracellular matrix protein. The matrix location of MFS mutations means even cells not expressing the mutated gene may directly interact with the diseased protein product, posing an exciting test case for single cell proteomics. As such, we have described the effects of sex differences and aneurysm pathology on these single cell proteomes to demonstrate how this new technology can be used to uncover changes in the abundance and diversity of major cell-type classifications. Specifically, we chose Marfan aortic root aneurysms as a pathological variation as it is known for its shift in subtypes of SMCs and has been extensively assayed through single cell RNA sequencing, providing us with an ability to directly compare the two single cell modalities head to head and combine them into one master signature^19^. While single cell RNA sequencing has some technical advantages, single cell proteomics, despite the challenges of sequencing peptides, provides a closer understanding of the true functional state of a cell.

We describe here a protocol that combines advances in microfluidic technology to allocate single cells into 384-well plates with additional adaptations in TOF mass spectrometry to sequence the proteome from a single cell. On average we were able to detect roughly 500 proteins per cell which represents about 15% of the proteome detected using bulk tissue proteomics on similar samples^32^. After initial cell clustering of each biological replicate, we observed considerable batch effect, not unlike other forms of large-scale genetic profiling, but we were able to correct for such differences to ensure that each replicate contributed uniformly to most cell clusters. Mouse aortic root tissue is extremely small starting material, yet we successfully profiled 3,745 cells with at least 200 proteins across 24 samples. These cells clustered using the leiden algorithm into 16 groups featuring protein markers that allowed identification of all known major cell types in aortic mouse tissue. This included several subtypes of abundant cells like SMCs and fibroblasts. SMCs were by far the most abundant and diverse cell type, as anticipated given the histological structure of the aorta and the relative pluripotency of SMCs. In fact, we identified a total of 7 SMC subtypes characterized by relatively high expression of contractile markers like Myh11, Smtn, Vcl, and Tagln. Nevertheless, there was considerable variation in the expression of these canonical markers throughout the different subtypes namely those with high expression in the SMC2, SMC3, and SM6 clusters versus low expression in the SMC1, SMC4, and SMC5 groups. One of the most well described variations in SMCs in vascular tissue divides this major cell type into two categories: those with a more quiescent and contractile like state and those with a more proliferative synthetic like phenotype^33^. The abundance of contractile markers detected in the single cell proteome could reflect these earlier classifications. The synthetic like clusters also had significantly more expression of the proteins Lrp1, Lig3, and Prss2. Lrp1 is a well characterized SMC receptor that plays a critical role in the integrity of vessel walls. Disruptions in the Lrp1 gene have been linked to both abdominal and thoracic aneurysms and may involve Lrp1’s ability to regulate TGFβ and proteases in the extracellular matrix. Interestingly, specific deletion of Lrp1 in SMCs results in ascending thoracic aortic aneurysms in mice, similar to Marfan’s syndrome^34^. Lig3 and Prss2, on the other hand, have known functionality in the metabolic and mitochondrial stability of SMCs, pathways also linked to aneurysm biology^34^. Two additional interesting findings included the adipocyte-like profile of SMC7 despite its extremely high expression of Myh11, and the unique expression of the capsaicin channel Trpv1 in the SMC5 cluster. RNA-seq analysis of certain types of adipose tissue has recently defined a group of Trpv1^+^ adipocyte progenitor cells that originate from vascular SMCs^35^. Therefore, it is possible that SMC5 and SMC7 could represent phases of a lineage transition between SMCs in the wall of the aorta and perivascular fat in the adventitia. Future experiments tracking these lineages in mouse models could uncover a novel understanding of the development of vasculature which may play a role in pathological states like Marfan syndrome.

Among the weaknesses of bulk proteomics is an inability to distinguish if DEPs represent changes within certain cells or changes in the abundance of cell types. One of the principal objectives of this study was to try to identify novel cellular specific differences between male and female mice and between wild-type and Marfan mice. In each comparison there were few proteins that met multiple comparison testing cutoffs to be considered Bonafide DEPs. This could be due to many reasons including more prominent differences in cell types as opposed to within cell type changes, and lack of power to overcome technical and/or biological variability. With regards to sex, there were more DEPs in fibroblasts, endothelial cells, and SMC groups, which could simply be due to these cohorts having more cells. There was one comparison that yielded a significantly higher number of DEPs than all others, namely the DEPs between wild-type and Marfan mice within endothelial cells from male mice only. These DEPs were significantly enriched in functions related to endothelial to mesenchymal transition. EndMT has been demonstrated as a potential driver of aneurysm pathogenesis in patients with bicuspid aortic valves^36^. Recent studies demonstrating the roles of miRNAs in Marfan aortopathy have shown a significant pattern of endMT in Marfan aortic aneurysms compared to other genetic forms of the disease. These findings suggest that increased endMT, through nitric oxide, TGFβ, and Wnt signaling, may destabilize the intima increasing the rate of aneurysm growth and dissection. Changes in endMT may also influence endothelial signaling to the SMCs in the medial layer, which likely are the cells most involved in the pathogenesis of aneurysm development^36,37^. The fact that the endMT pathway seems more disturbed in male mice is consistent with previous studies which have shown a sex related exaggeration in the aneurysm phenotype in both Marfan mouse models and human Marfan patients^15, 38^. Future studies are needed to clarify if these changes in endMT occur only during embryogenesis or continue to happen during post-embryologic development.

By combining single cell informatics from multiple omic platforms, we can begin to build genetic networks on the cellular level with the hope of uncovering novel pathways within and between cells that contribute to aneurysm pathogenesis. As a result, we took advantage of the publicly available single cell RNA-seq data performed on the same Marfan mouse model used in our study. These RNA studies were fortunately performed on similar aortic tissue at a similar stage of development^19^. Interestingly, both trans-analyte and integrated analyses revealed that outside the overlap in major cell types, there was very little similarity between the two omic modalities. There are several possible explanations for these findings including: differences in the genetic background strain of the *Fbn1^C1041G^* mutants (C57Black6 for scTranscriptome and 129 for scProteome), the relatively large number of RNA features (1000s) compared to protein features (100s), the fact that RNA represents more active gene transcription while protein could be more reflective of a functional steady state, different sources of technical and biological variation across studies that were unaccounted for in our models, and underdevelopment of dedicated computational programs that can integrate single cell omics, particularly proteomics data. Despite this poor overlap, the LIGER integrated SMC projection from the single cell proteomic and transcriptomic datasets did identify unique clusters of multi-omic-defined cells, some of which demonstrated altered proportions across wild-type and Marfan mice. In particular, key functional components of the contractile apparatus were enriched in multi-omic clusters with higher contributions from wild-type cells, which highlights the importance of contractility and its regulation in maintaining vessel wall integrity, while integrated leiden clusters with significant contributions from Marfan cells contained genes related to angiotensin, TGFβ, and protease signaling. It is particularly interesting that Lrp1 and Ace were two multi-omic features that uniquely defined MFS enriched cell types, because these two genes have known contributions to the Marfan phenotype^34, 39^. While traditionally considered an endothelial-enriched protein, expression of Ace specifically on medial SMCs has recently been mechanistically linked to atherosclerotic propensity in mouse models, suggesting an important role for this protein in SMC biology and this warrants further mechanistic investigation in aneurysm^40^. Also of interest, Tpm4 was a marker alongside Ace in MFS enriched cluster 4, and this gene has been highlighted as upregulated during SMC dedifferentiation and is enriched in atherosclerotic plaque SMCs^41^. Taken together, from a multiomic analysis it is clear that cluster 4 is transcribing, translating, and stably expressing higher levels of markers for SMC dedifferentiation. These markers have been linked previously to other aortic pathologies, but our data are the first to our knowledge to implicate them in MFS aneurysm.

In summary, this single cell proteomic analysis, the first of its kind, has revealed novel insights into potential MFS specific cellular phenotypes, including suggesting the involvement of endMT in altered endothelial function in MFS, as well as potentially key roles for SMC Ace and Lrp1. With improvements in both forms of single cell technology and advances in other forms of single cell omics such as metabolomics, lipidomics, noncoding RNA omics, and epigenomics, we hope to fully map all pertinent signaling pathways that contribute to the pathology of thoracic aneurysms, and together with the findings already reported here, leverage these valuable insights toward developing an effective pharmacological intervention to allay aneurysm progression in these patients.

## Acknowledgements

We would like to thank and acknowledge the Cedars Sinai Proteomics and Metabolomics Core facility for their effort in generating these data. We also thank the Fischbein laboratory for the deposition of and public access to the single cell transcriptomics data.

## References

1. J. Lim, C. Park, M. Kim, H. Kim, J. Kim and D. S. Lee, Exp Mol Med 56 (3), 515–526 (2024).

2. A. Ghazalpour, B. Bennett, V. A. Petyuk, L. Orozco, R. Hagopian, I. N. Mungrue, C. R. Farber, J. Sinsheimer, H. M. Kang, N. Furlotte, C. C. Park, P. Z. Wen, H. Brewer, K. Weitz, D. G. Camp, 2nd, C. Pan, R. Yordanova, I. Neuhaus, C. Tilford, N. Siemers, P. Gargalovic, E. Eskin, T. Kirchgessner, D. J. Smith, R. D. Smith and A. J. Lusis, PLoS Genet 7 (6), e1001393 (2011).

3. C. Vogel and E. M. Marcotte, Nat Rev Genet 13 (4), 227–232 (2012).

4. M. Yang, F. Petralia, Z. Li, H. Li, W. Ma, X. Song, S. Kim, H. Lee, H. Yu, B. Lee, S. Bae, E. Heo, J. Kaczmarczyk, P. Stępniak, M. Warchoł, T. Yu, A. P. Calinawan, P. C. Boutros, S. H. Payne, B. Reva, E. Boja, H. Rodriguez, G. Stolovitzky, Y. Guan, J. Kang, P. Wang, D. Fenyö and J. Saez-Rodriguez, Cell Syst 11 (2), 186–195.e189 (2020).

5. H. Srivastava, M. J. Lippincott, J. Currie, R. Canfield, M. P. Y. Lam and E. Lau, PLoS Comput Biol 18 (11), e1010702 (2022).

6. X. Cheng, K. Wang, Y. Zhao and K. Wang, Cell Death Discovery 9 (1), 275 (2023).

7. V. Petrosius and E. M. Schoof, Translational Oncology 27, 101556 (2023).

8. L. Sun, K. M. Dubiak, E. H. Peuchen, Z. Zhang, G. Zhu, P. W. Huber and N. J. Dovichi, Anal Chem 88 (13), 6653–6657 (2016).

9. B. Budnik, E. Levy, G. Harmange and N. Slavov, Genome Biology 19 (1), 161 (2018).

10. A. D. Brunner, M. Thielert, C. Vasilopoulou, C. Ammar, F. Coscia, A. Mund, O. B. Hoerning, N. Bache, A. Apalategui, M. Lubeck, S. Richter, D. S. Fischer, O. Raether, M. A. Park, F. Meier, F. J. Theis and M. Mann, Mol Syst Biol 18 (3), e10798 (2022).

11. S. Kreimer, A. Binek, B. Chazarin, J. H. Cho, A. Haghani, A. Hutton, E. Marbán, M. Mastali, J. G. Meyer, T. Mesquita, Y. Song, J. Van Eyk and S. Parker, Anal Chem 95 (24), 9145–9150 (2023).

12. C. Ctortecka, N. M. Clark, B. Boyle, A. Seth, D. R. Mani, N. D. Udeshi and S. A. Carr, bioRxiv (2024).

13. K. A. Groth, K. Stochholm, H. Hove, K. Kyhl, P. A. Gregersen, N. Vejlstrup, J. R. Østergaard, C. H. Gravholt and N. H. Andersen, Clin Res Cardiol 106 (2), 105–112 (2017).

14. L. F. Hiratzka, G. L. Bakris, J. A. Beckman, R. M. Bersin, V. F. Carr, D. E. Casey, Jr., K. A. Eagle, L. K. Hermann, E. M. Isselbacher, E. A. Kazerooni, N. T. Kouchoukos, B. W. Lytle, D. M. Milewicz, D. L. Reich, S. Sen, J. A. Shinn, L. G. Svensson and D. M. Williams, J Am Coll Cardiol 55 (14), e27–e129 (2010).

15. L. Saddic, S. Escopete, L. Zilberberg, S. Kalsow, D. Gupta, M. Eghbali and S. Parker, Int J Mol Sci 24 (17) (2023).

16. X. Chen, D. L. Rateri, D. A. Howatt, A. Balakrishnan, J. J. Moorleghen, L. A. Cassis and A. Daugherty, PLoS One 11 (4), e0153811 (2016).

17. H. Wei, J. H. Hu, S. N. Angelov, K. Fox, J. Yan, R. Enstrom, A. Smith and D. A. Dichek, J Am Heart Assoc 6 (1) (2017).

18. E. G. MacFarlane, S. J. Parker, J. Y. Shin, B. E. Kang, S. G. Ziegler, T. J. Creamer, R. Bagirzadeh, D. Bedja, Y. Chen, J. F. Calderon, K. Weissler, P. A. Frischmeyer-Guerrerio, M. E. Lindsay, J. P. Habashi and H. C. Dietz, The Journal of Clinical Investigation 129 (2), 659–675 (2019).

19. A. J. Pedroza, A. R. Dalal, R. Shad, N. Yokoyama, K. Nakamura, P. Cheng, R. C. Wirka, O. Mitchel, M. Baiocchi, W. Hiesinger, T. Quertermous and M. P. Fischbein, Arterioscler Thromb Vasc Biol 42 (9), 1154–1168 (2022).

20. A. J. Pedroza, Y. Tashima, R. Shad, P. Cheng, R. Wirka, S. Churovich, K. Nakamura, N. Yokoyama, J. Z. Cui, C. Iosef, W. Hiesinger, T. Quertermous and M. P. Fischbein, Arterioscler Thromb Vasc Biol 40 (9), 2195–2211 (2020).

21. A. T. Kong, F. V. Leprevost, D. M. Avtonomov, D. Mellacheruvu and A. I. Nesvizhskii, Nat Methods 14 (5), 513–520 (2017).

22. F. A. Wolf, P. Angerer and F. J. Theis, Genome Biology 19 (1), 15 (2018).

23. V. Demichev, C. B. Messner, S. I. Vernardis, K. S. Lilley and M. Ralser, Nat Methods 17 (1), 41–44 (2020).

24. A. Behdenna, M. Colange, J. Haziza, A. Gema, G. Appé, C.-A. Azencott and A. Nordor, BMC Bioinformatics 24 (1), 459 (2023).

25. A. Butler, P. Hoffman, P. Smibert, E. Papalexi and R. Satija, Nat Biotechnol 36 (5), 411–420 (2018).

26. T. Stuart, A. Butler, P. Hoffman, C. Hafemeister, E. Papalexi, W. M. Mauck, 3rd, Y. Hao, M. Stoeckius, P. Smibert and R. Satija, Cell 177 (7), 1888–1902.e1821 (2019).

27. J. D. Welch, V. Kozareva, A. Ferreira, C. Vanderburg, C. Martin and E. Z. Macosko, Cell 177 (7), 1873–1887.e1817 (2019).

28. J. Liu, C. Gao, J. Sodicoff, V. Kozareva, E. Z. Macosko and J. D. Welch, Nat Protoc 15 (11), 3632–3662 (2020).

29. M. Kanehisa and S. Goto, Nucleic Acids Res 28 (1), 27–30 (2000).

30. E. Y. Chen, C. M. Tan, Y. Kou, Q. Duan, Z. Wang, G. V. Meirelles, N. R. Clark and A. Ma’ayan, BMC Bioinformatics 14, 128 (2013).

31. Y. Hao, T. Stuart, M. H. Kowalski, S. Choudhary, P. Hoffman, A. Hartman, A. Srivastava, G. Molla, S. Madad, C. Fernandez-Granda and R. Satija, Nature Biotechnology 42 (2), 293–304 (2024).

32. S. J. Parker, A. Stotland, E. MacFarlane, N. Wilson, A. Orosco, V. Venkatraman, K. Madrid, R. Gottlieb, H. C. Dietz and J. E. Van Eyk, Am J Physiol Heart Circ Physiol 315 (5), H1112–h1126 (2018).

33. S. S. Rensen, P. A. Doevendans and G. J. van Eys, Neth Heart J 15 (3), 100–108 (2007).

34. P. Boucher, W. P. Li, R. L. Matz, Y. Takayama, J. Auwerx, R. G. Anderson and J. Herz, PLoS One 2 (5), e448 (2007).

35. F. Shamsi, M. Piper, L.-L. Ho, T. L. Huang, A. Gupta, A. Streets, M. D. Lynes and Y.-H. Tseng, Nature Metabolism 3 (4), 485–495 (2021).

36. S. Maleki, F. A. Poujade, O. Bergman, J. R. Gådin, N. Simon, K. Lång, A. Franco-Cereceda, S. C. Body, H. M. Björck and P. Eriksson, Front Cardiovasc Med 6, 182 (2019).

37. S. Terriaca, M. G. Scioli, F. Bertoldo, C. Pisano, P. Nardi, C. R. Balistreri, D. Magro, B. Belmonte, L. Savino, A. Ferlosio and A. Orlandi, Cells 13 (15) (2024).

38. K. J. Grubb and I. L. Kron, Semin Thorac Cardiovasc Surg 23 (2), 124–125 (2011).

39. C. Yu and R. W. Jeremy, Int J Cardiol Heart Vasc 18, 71–80 (2018).

40. X. Chen, D. A. Howatt, A. Balakrishnan, J. J. Moorleghen, C. Wu, L. A. Cassis, A. Daugherty and H. Lu, Arterioscler Thromb Vasc Biol 36 (6), 1085–1089 (2016).

41. M. Abouhamed, S. Reichenberg, H. Robenek and G. Plenz, Eur J Cell Biol 82 (9), 473–482 (2003).

